# Wheel running, but not home cage activity in Digital Ventilated Cages® is impaired in a mouse model of breast cancer-induced bone pain

**DOI:** 10.1101/2024.11.13.623396

**Authors:** Chelsea Hopkins, Ida Buur Kanneworff, Birgitte Rahbek Kornum, Anne-Marie Heegaard

## Abstract

**Introduction:** Cancer-induced bone pain (CIBP) due to metastatic breast cancer is common and debilitating. Effective, long-term treatment options have side effects that reduce patients’ quality of life. Preclinical models are a valuable tool to test novel analgesics, but new methods that are translationally and clinically relevant are necessary. This study aimed to assess spontaneous pain-like behaviour of home cage activity and wheel running in Digital Ventilated Cages®.

**Method:** Twenty BALB/cAnNHsd mice were housed in Digital Ventilated Cages® from Tecniplast® with GYM500 home cage running wheels. Mice were assessed by limb use analysis and static weight bearing to determine the development of CIBP and this was compared to the dark-phase home cage activity and wheel running in the Digital Ventilated Cages®. Ten mice underwent 4T1-Luc2 mammary gland adenocarcinoma cell inoculation into the right femur to establish CIBP and another ten mice underwent a sham surgical procedure.

**Results:** The 4T1-inoculated mice displayed pain-like behaviour in limb use and weight bearing tests, demonstrating a preference for the contralateral limb. The limb use scores were compared with home cage activity and wheel running. A reduced wheel running distance corresponded to reduced limb use scores, with the shortest wheel running distances corresponding to the lowest limb use scores. However, this behavioural pattern was not observed in home cage activity, which remained consistent over the study period.

**Discussion:** Reduced wheel running corresponded with reduced limb use scores. This suggests that wheel running behaviour is affected by the development of metastatic breast cancer. However, the home cage activity was not influenced by disease development. Digital Ventilated Cage® wheel running may be a useful behavioural assessment of spontaneous pain-like behaviour of CIBP and may be useful to assess analgesic efficacy.

## Introduction

In patients with advanced breast cancer, 58% will experience bone metastases [1] and of those, 68% will experience bone pain [2]. These patients commonly experience worse pain and quality of life than patients with metastases in non-bone tissue. It has also been reported that of the patients with painful bone metastases, 97% are on analgesics; 55% taking opioids and 42% taking non-opioids [2]. However, long-term pain relief therapies (e.g. non-steroidal anti-inflammatory drugs, opioids) lead to an increase in side effects that impair quality of life [3]. Preclinical *in vivo* models are an essential tool to investigate nociceptive mechanisms and novel analgesics [4].

Evoked pain tests are common in pre-clinical settings to assess mechanical hypersensitivity (i.e. Von Frey test) and thermal hypersensitivity [5, 6]. However, spontaneous, ongoing pain is considered a more relevant issue for patients, and this is difficult to test *in vivo* [7]. Pain is not only a sensory condition, but may also encompass stress, anxiety, depression, and limit social interactions. The sensory and affective conditions should be assessed in combination to develop a well-rounded model of pain. Furthermore, *in vivo* tests often occur during the day over a short period, when rodents are least active, which means that a measure of spontaneous pain fluctuations may be difficult to gauge [8].

Tecniplast SpA has developed Digital Ventilated Cages® (DVCs®) that can track home cage activity and monitor wheel running. The DVC® rack is equipped with an electromagnetic sensing board with capacitance sensing technology to assess home cage activity; the board contains 12 electrodes that emit an electromagnetic field and when a mouse moves over these fields, it creates a disturbance that is recorded and measured. Readings are obtained continuously and unobtrusively when the cages are in the rack, allowing for continuous monitoring without handling interference during the 12-hour dark phase when mice are more active [9]. This provides the following benefits: evaluate ethologically relevant behaviour in a familiar home cage environment, minimise researcher interference, and reduce handling stress, thus mitigating common confounding factors in pain behavioural studies. DVCs have been used in previous studies to assess circadian rhythms [10-12], effect of standard procedures on cage activity [12, 13], severity monitoring [13, 14], and recently a novel treatment for osteoarthritis pain (rest disturbance metric only) [15]. Wheels with perpendicular magnets have been developed that can be placed in the cages, and their rotation, speed, and distance monitored automatically. However, wheel running in DVCs® has never been established or measured as a behavioural outcome.

A distinction should be made between home cage activity, horizontal activity, and vertical activity. Home cage activity describes the movement and activity around the home cage environment where a rodent is permanently housed. Horizontal activity describes movement in a two-dimensional testing area where a rodent is allowed to explore freely for a specified time. Vertical activity may be described as rearing, where the rodent stands only on its hind legs in order to extend vertically upwards. Reduced horizontal activity has been observed in fracture [16], arthritic [17], inflammatory [18, 19], neuropathic [19], and cancer-induced bone pain models [20]. These experiments were conducted over short (60-120 min) [17-19] and 20-hour periods [16, 20], but not in standard home cages. These experimental setups may introduce confounding stress and exploratory behaviours, and they require additional time to set up and carry out the experiment. Urban et al. [21] used a home cage monitoring system to demonstrate that a neuropathic pain model resulted in reduced cage activity in a neuropathic pain model in a BALB/c mouse strain compared to naïve mice. The system is similar to PhenoTyper® developed by Noldus®, which uses LED units and cameras to measure behaviour [22]. Wheel running (during a short period) has been used in previous studies to assess the development of pain-like behaviour in models of prostate cancer-induced bone pain [23], inflammation, and neuropathic pain [18, 19, 24].

The aim of this study is to determine if home cage activity and wheel running are affected by the development of breast CIBP. These metrics are compared to limb use behaviour, an established test of pain-like behaviour in this mouse model.

## Method

### Animals

Twenty 5-week-old female BALB/cAnNHsd mice were used in this study; ten mice were inoculated with 4T1-Luc2 cells and ten mice underwent sham surgery. All experiments were conducted according to the Danish Animal Experiments Inspectorate (Copenhagen, Denmark, 2020_15_0201_00439). Mice were purchased from Envigo (USA) and housed in a certified specific pathogen-free facility at the University of Copenhagen where all experiments were conducted. The research unit maintains a temperature of 22±2°C and humidity of 55±10% in housing and experimental rooms. The light/dark cycle was 12-hours and light was kept at 60% intensity (7am-7pm). During the acclimatisation period, mice were housed in GM500 individually ventilated cages with GYM500 cage wheels (Tecniplast®, Italy) and during the testing period, mice were housed in the DVCs® within the DVC® rack (Tecniplast®, Italy). All cages contained wood-chip bedding material (Tapvei 2HV, Brogaarden, Denmark), one transparent red housing unit (Polycarbonate Mouse Tunnel, Datesand, UK), a running wheel (DVC® GYM500, Tecniplast®, Italy), nesting material (paper shavings, Brogaarden, Denmark), and a wooden gnawing block (Aspe små klodser, Brogaarden, Denmark). Standard chow (Altromin 1324, Brogaarden, Denmark) and tap water were provided ad libitum. Fresh food and water was provided once per week and cages were changed every two weeks. Severity monitoring was conducted during the baseline experiments, D1-D4 after surgery, and every experimental day thereafter. Severity monitoring included weight loss/gain, coat condition, aggression, mobility, etc. A predefined scoring system was implemented to ensure that mice did not surpass the humane endpoint. The humane endpoint was a mouse reaching a pre-defined severity score of six or a limb use score zero. The experimental endpoint was defined as a 4T1-inoculated mouse reaching a limb use score of zero and control mice reached the experimental endpoint when all 4T1-inoculated mice had been euthanised. Mice were only handled using the tunnel handling method or scooping them with hands to minimise stress. All mice were euthanised by cervical dislocation.

### Study Design

Mice were stratified into the 4T1-inoculated group and sham control group according to weight-bearing baseline data and body weight by the primary researcher (CH) to ensure that both groups had an equal average baseline behaviour and weight. The experimenter (IBK) was blinded throughout the experiment to the group allocation. Blinding was not possible during data analysis due to the nature of the experiment where group allocation is revealed during the course of the study. Each mouse acted as one experimental unit. Mice were excluded from the experiment if they demonstrated any indication of poor health prior to surgery, if they did not completely heal after surgery prior to D5, if they demonstrated symptoms of tendon displacement (e.g. limping, unable to extend ipsilateral limb), or if they did not run on their cage wheel. All behavioural experiments that were conducted outside the cage were done in the morning (9am-12pm) at the same time for each mouse.

### Experimental Timeline

Figure 1 demonstrates the timeline of the full experiment. Mice were introduced to the facilities at 5-weeks old. They acclimatised for one week prior to baseline experiments without interruption. At D-7 they were transferred to individually housed DVCs®. From D-7 to D-5, mice were trained and habituated to the static weight bearing procedure. On D-4, no behavioural or experimental procedures were conducted prior to the dark phase to obtain cage activity and wheel running data that is not influenced by experimenter handling. On D-3 and D-2, mice underwent limb use and weight bearing baseline tests. The following dark phases were used as baseline assessments to determine the effect of handling on cage activity and wheel running. Baseline x-ray images were collected on D-1 and surgery was conducted on D0. Post-surgical care was conducted from D1-D4.

**Figure 1:**
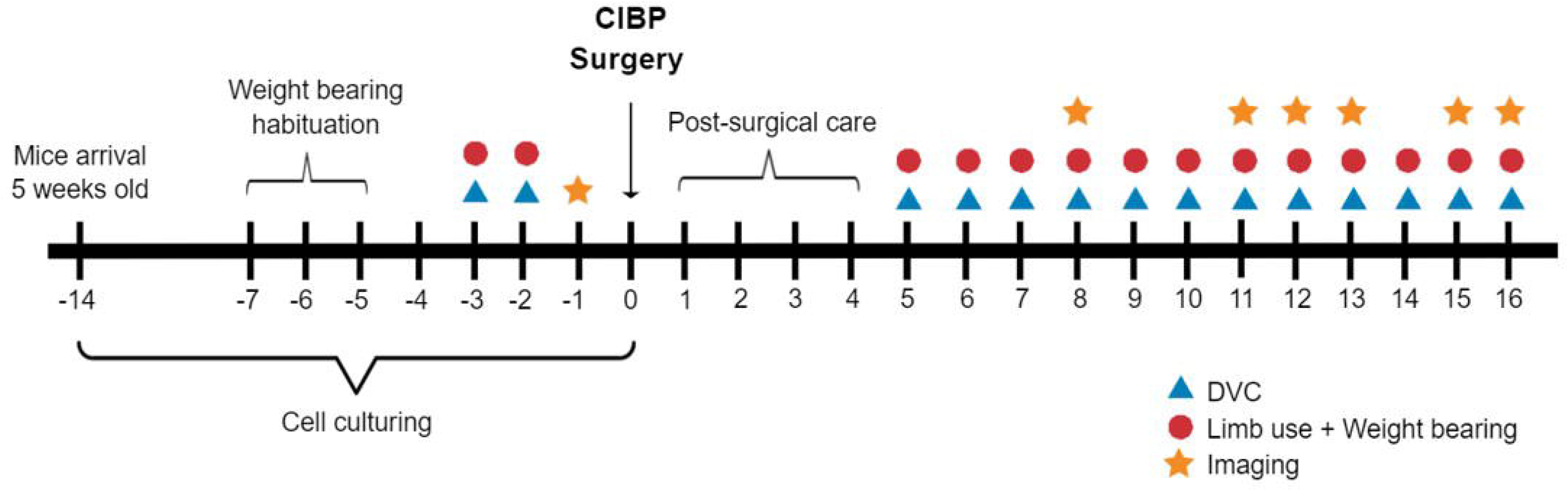
Experimental timeline demonstrating animal behaviour and cell culture procedures over the course of the experiment. (CIBP: cancer-induced bone pain, DVC: Digital Ventilated Cage®)

From D5 onwards, limb use and weight bearing were conducted every day during the light phase and cage activity and wheel running were only assessed in the dark phase. Imaging was conducted on D8 and prior to euthanasia when mice reached the experimental or humane endpoint.

### Cell Culture

Mouse mammary gland adenocarcinoma cells (Bioware Ultra Cell Line 4T1-luc2; Caliper Life Sciences, Teralfene, Belgium; ATCC (CRL-2539^™^) parental line) were cultured as previously described [25, 26]. Cells were cultured in vented sterile cell culture flasks (75cm^2^; Cellstar®, Greiner, Austria) in a sterile environment. Cells were maintained in RPMI 1640 cell culture media without phenol red (Gibco®, USA), supplemented with 10% heat-inactivated foetal bovine serum (Gibco®, USA) and 1% penicillin-streptomycin-glutamine (Gibco®, USA). Prior to surgery, cells were cultured for two weeks and then split two days before to ensure log-phase growth. To split and harvest the cells, 0.5% trypsin-EDTA (Gibco®, USA) was applied to the cells for 5 minutes. Trypsin was inactivated with regular cell media, the solution was centrifuged, supernatant discarded, and cells resuspended in Hank Balanced Salt Solution (HBSS; Gibco®, USA) to a concentration of 10^6^ cells/ml. Cells were kept on ice thereafter.

### Cancer Cell Inoculation in Femur

Surgery was conducted as previously described, with minor modification [26]. Mice were anaesthetised with an intraperitoneal xylazine/ketamine cocktail (43 mg/kg ketamine, MSD Animal Health, AN Boxmeer, the Netherlands; 6mg/kg xylazine, Rompun vet, Bayer, Germany). Mice were maintained on 1-1.2% isoflurane (1000mg/g isoflurane, Attane vet, ScanVet, UK) throughout surgery. Eye ointment (Ophtha A/S, Activis Group, Gentofte, Denmark) was applied to the eyes, 0.9% saline was injected subcutaneously to prevent post-surgical dehydration, and 5 mg/kg carprofen (Carprosan Vet, Dechra, the Netherlands) was injected subcutaneously. Mice were placed on a heated surgery table in a supine position. A small incision was made over the patella tendon and in the connective tissue on the medial side of the tendon. The tendon was positioned to the lateral side of the knee. A 30-G needle was used to manually drill a hole into the femoral epiphysis until the medullary cavity was reached. Using a 0.3 ml insulin syringe (BD, USA), 10 μl of 4T1-luc2 cells (4T1-inoculated group) or HBSS was injected into the cavity and incubated for 1 minute. The hole was filled with bone wax (Harvard Apparatus, USA) and the patella tendon moved back to the central position. The region was thoroughly rinsed with 0.9% saline and the incision closed with two medical clips (Michel Suture Clips, Agnthos, Sweden). Mice were maintained in a prone position on a heat map until they regained mobility. On D1, mice were administered 5 mg/kg carprofen subcutaneously for post-surgical analgesia.

### Digital Ventilated Cage® - Home Cage Activity and Wheel Running

The electromagnetic wheels (GYM500, Tecniplast, Italy) were secured to the inner side of the DVCs®. When the wheel spins, the electromagnetic field produced by the magnets can be detected and tracked. The cumulative distance during the dark phase was assessed. Mice are acclimatised for one week in group housing (5 mice per cage) with a cage wheel in the Individual Ventilated Cages®. This is an essential training and habituation step for the mice. Mice without wheel habituation may not use the running wheel during the testing period. After the acclimatisation period, mice were single housed in the DVCs® and rack. Assessments conducted outside of the cages were conducted during the same time in the morning, for the same period. This arrangement was practiced to ensure that dark phase behaviour was not influenced by behaviour experiments and handling during the light phase. When mice were removed from their cages during the behavioural experiments, bedding material was moved away from the wheel; bedding can interfere with the wheel mechanism and magnets, preventing accurate readings. The rack was placed in the housing room away from the entrance door to prevent excess interference and in a position with minimal human traffic. The mice were housed with half the mice from each group at the top and bottom of the rack to determine if position in the rack was a confounding factor. Analysis was conducted on data collected from the dark phase (7pm-7am). Mice with a peak hourly distance of less than 500 m were excluded from the experiment.

Home cage activity metrics and the associated calculations have been described previously [9] and shown to be comparable to manually analysed video analysis of mouse behaviour in the cage. When the emitted electromagnetic field is disturbed, it will be detected and recorded. The calculated activity density is calculated every minute (four recordings per minute) and the average home cage activity per hour was assessed.

When comparing data from different tests, the cage activity and wheel running data collected from the dark phase prior to the limb use and weight bearing tests were matched.

### Limb Use

Mice were placed in an empty transparent box for ten minutes to habituate. Thereafter, mice were transferred to an identical testing arena where they were monitored for three minutes. Mice were scored according to the following scale:

4 – Normal gait

3 – Insignificant limping

2 – Significant limping and shift in bodyweight towards the healthy limb

1 – Significant limping and partial lack of use of the ipsilateral leg

0 – Total lack of use of the ipsilateral leg

Different boxes were used for each mouse.

### Static Weight Bearing

Static weight bearing was assessed using the Incapacitance Tester (Version 5.2, Linton, UK). Mice were habituated and trained to maintain a testing position (hind paws individually placed on two weight plates) prior to baseline measurements. Triplicate measures of 3-second readings were obtained for every mouse and the ratio of weight distribution was calculated.

### X-ray and Bioluminescent Imaging

Mice underwent imaging on D8 and when the mice reached their experimental or humane endpoint. Mice were injected intraperitoneally with 150mg/kg D-Luciferin (PerkinElmer, Inc., USA) nine minutes prior to imaging. Mice were sedated with 2.5% isoflurane (1000 mg/g isoflurane, Attane vet, ScanVet, UK) and maintained throughout the imaging procedure. Lumina XR apparatus (Caliper Life Sciences, Teralfene, Belgium) was used to obtain both x-ray and bioluminescent images in triplicate for 4T1-inoculated mice. One x-ray image was obtained for the sham control mice.

### Statistics

The *a priori* sample size calculation was based on previous studies conducted on the same model, with power set at 80% and statistical significance set at p<0.05. Calculated group size was increased by 20% to account for possible exclusion.

Data collected by the DVC® system was extracted with data summarised for every hour. Home cage activity data was presented as previously described [9]. Wheel running was presented as cumulative running distance per hour.

Due to the experimental design of mice euthanised during the course of the study, the last observation was carried forward to account for missing measures at later time points.

All graphs and plots presented were generated with GraphPad Prism 9.3.1 (GraphPad Software, United States). The parametric data (weight bearing, home cage activity, wheel running) was analysed using the mixed-effects model, one-way ANOVA, or two-way ANOVA with appropriate corrections for multiple comparison (see specific information in figure legends) in GraphPad Prism. The non-parametric data (limb use) was analysed by the Friedman test in SAS® 9.4 (SAS Institute, United States) with pairwise Mann-Whitney U tests in Graphpad Prism.

## Results

Mice in the sham group did not demonstrate significantly decreased behaviour in any of the tests over the course of the experiment. Mice that were inoculated with 4T1-Luc2 cells in the right femur were evaluated daily by activity monitoring in DVCs®, limb use, and weight bearing. Presence of pain-like behaviour in the 4T1-inoculated mice was confirmed by a significant decrease in limb use score compared to baseline from D12 (Figure 2A) and weight bearing ratio from D10 (Figure 2B) after inoculation.

**Figure 2:**
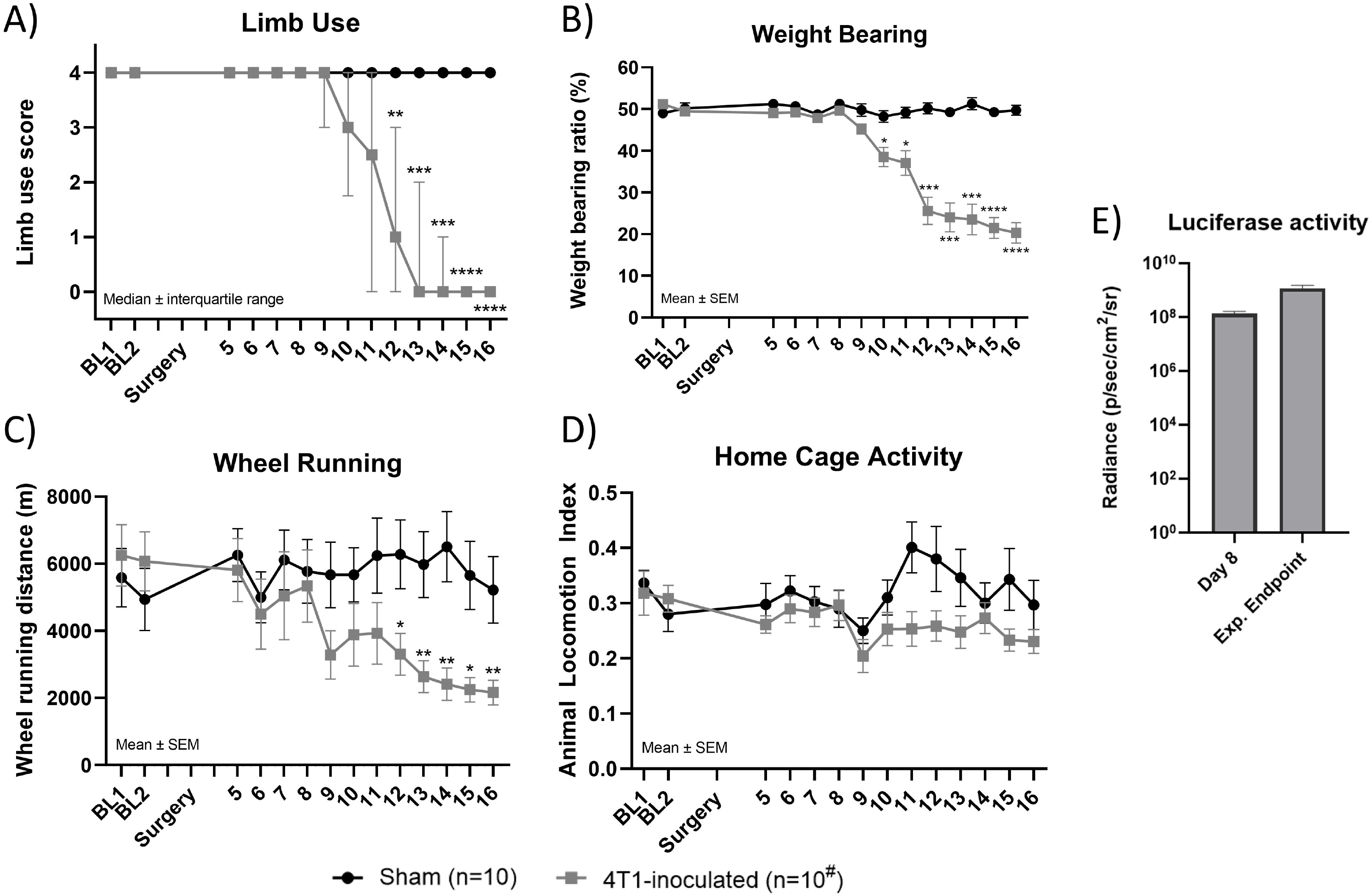
Effect of intrafemoral inoculation of 4T1-Luc2 cells on A) limb use, B) weight bearing, C) wheel running, D) home cage activity, E) luciferase activity. Cumulative wheel running distance and average animal locomotion index data is hourly DVC®-data from the dark phase (7pm-7am) preceding a study day. Statistics are performed with (A) Friedman test with pairwise Mann-Whitney U tests or (B, C, D, E) mixed-effects model with Sidak multiple comparisons test. ^#^Data analyses at last observation carried forward to account for animals euthanised during the study and comparisons were made to BL. (BL: baseline; *: p<0.05; **: p<0.01; ***: p<0.001, ****: p<0.0001; ^#^: last observation carried forward)

4T1-inoculated mice showed significantly decreased wheel running from D12 (Figure 2C) compared to baseline. Horizontal activity did not change within the experimental timeframe (Figure 2D). Tumour growth in the ipsilateral femur was confirmed by *in vivo* luciferase imaging on D8 and upon reaching the humane and experimental endpoint (Figure 2E), which showed that all 4T1-inoculated mice that developed pain-like behaviour, developed a tumour in their inoculated limb.

To evaluate the relationship between activity measures and an established measure of pain-like behaviour, the DVC®-data from the dark phases were matched to the limb use score obtained the following study day (Figure 3A, 3B). 4T1-inoculated mice reaching a limb use score of 2 (significant limping) or 0 (complete lack of use of the ipsilateral limb) showed significantly lower wheel running distance compared to baseline throughout the whole dark phase (Figure 3A, 3C). The horizontal activity did not differ between any limb use scores (Figure 3B). Sham mice assessed during days that corresponded to a mean limb use score of 4, 2, and 0 in the 4T1-inoculated sham group were plotted, demonstrating no significant change in wheel running over time (Figure 3D).

**Figure 3:**
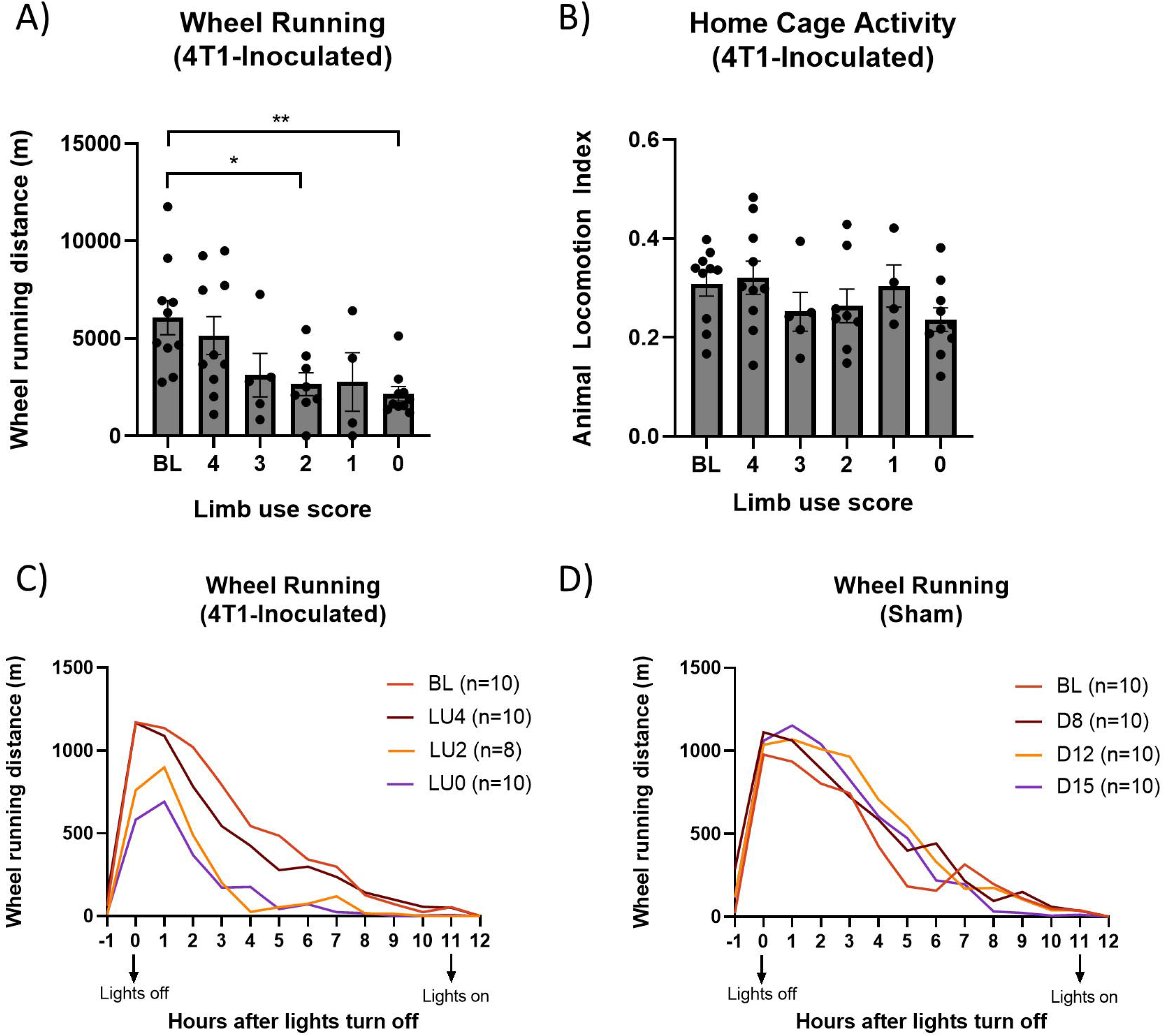
Relationship between limb use score and wheel running. A) Cumulative wheel running distance and B) home cage activity in 4T1-inoculated mice during the dark phase preceding the day the limb use was acquired for each mouse. C) Diurnal pattern of wheel running in 4T1-inoculated mice during the dark phase preceding the indicated limb use score. D) Diurnal pattern of wheel running in sham mice during the dark phase on days corresponding to an average limb use score of 4, 2, 0 within the 4T1-inoculated group. Statistics are performed with one-way ANOVA with Tukey’s test for multiple comparisons. Data is shown as mean ± SEM (A, B) and mean (C, D).

## Discussion

For the first time, DVCs® were used to assess spontaneous pain-like behaviour in a mouse model of breast CIBP. Wheel running was significantly reduced over time and reduced in concert with the reduction of limb use and static weight bearing. However, horizontal cage activity did not demonstrate a significant change over time or show any relationship with limb use or weight bearing.

Three measures of spontaneous pain-like development were reduced in this study as CIBP developed: limb use, weight bearing, and wheel running. Limb use and weight bearing are non-evoked behaviours, but they are performed during the light phase when mice are less active over a short period of time (total of 15 minutes to perform both tests). Previous studies on this model, using the same techniques have demonstrated that a decrease in limb use can be reversed with 10mg/kg morphine [27]. The reduction of wheel running occurred in parallel with the reduction of the limb use score. The first significant difference from baseline was observed on D12 in both limb use and wheel running (Figure 2A and 2C); also, wheel running significantly decreased comparably with limb use scores, demonstrating lower wheel running when mice scored lower in gait analysis. This suggests that wheel running is affected by the development of a malignant tumour and the associated nociceptive pathology. However, wheel running within this system holds the significant advantage of occurring during the more active dark phase, over the entire 12-hour period.

Wheel running has previously been performed in other models [18, 19, 23, 24]. However, these tests were performed during the light hours over a short period. Even so, similar to previous studies on prostate CIBP [23], inflammatory [18, 19], and chronic constriction injury models [19], this model also demonstrated significantly reduced wheel running behaviour. However, there are no previous studies demonstrating long-period wheel running and not in a home cage environment; this data suggests that this is a robust system capable of detecting the development of pain-like activity. There is evidence to suggest that these tests may be sex [24] and model [19] dependent. Care should be taken to understand the sex and strain constraints in this system, prior to implementing further studies. There is a legitimate concern that wheel running constitutes a stereotypic behaviour, defined as a repetitive behaviour with no goal or function [28] and it is to be avoided. However, a previous study demonstrates that wheel running is performed voluntarily by wild mice [29] and may be a rewarding and motivating activity [30].

In this study, there was no effect of CIBP on home cage activity. The only other comparable study on home cage systems was performed by Urban et al. [21] where a reduction of home cage activity was only observed in BALB/c mice with a chronic constriction injury. Home cage activity was not impacted in an inflammatory or spared nerve injury model [21]. Horizontal activity may be considered similar to home cage activity, which was affected in several models of bone pain [16-20]. However, home cage assessment offers an advantage of assessing pain-like behaviour over an extended period, instead of in an experimental system confounded by external factors. Again, sex and model within the DVC® system should be taken into account prior to using the system for further studies.

Analgesics are necessary to assess whether change of behaviour is due to pain-like development or other factors associated with stress or illness. However, spontaneous behaviour is constrained by the use of analgesics due the effect on overall behaviour. For example, previous studies on burrowing (also a spontaneous behaviour) demonstrated that most analgesics adversely affect the procedure. High doses of opioids (e.g. 10-30mg/kg morphine) are necessary to treat mouse models of CIBP, as weak analgesics are ineffective [27, 31]. Cobos et al. [18] have tested the use of analgesics in short-term wheel running assessments. In sham mice, a low dose of morphine (5mg/kg) significantly reduced wheel running. In mice with inflammatory-like pain, wheel running is partially reduce, but not significantly so, suggesting that morphine moderately restores wheel running behaviour in a pain-like model. This suggests that at least low doses of morphine could be used in intervention studies using wheel running as a behavioural outcome. However, an analgesic with a long half-life is required to span the 12-hour dark phase, or at least in this model, the first four hours when mice are most active. Morphine has a half-life of approximately 30 minutes, which would not work in this study design [32, 33], but buprenorphine hydrochloride with a 3-5 hour half-life or sustained release buprenorphine which lasts longer, may be more suitable alternatives [34, 35].

DVCs® also offer an advantage over other commercially available home cage systems. If Tecniplast® Individually Ventilated Cages are being used in a facility, DVCs® can be integrated easily, because their service and cleaning is similar to individually ventilated cages. They do not take up a lot of space, allowing 64 cages to be placed in a vertical rack, contrary to other systems that are larger and require more space for fewer cages. However, DVCs® are expensive (not compared to other home cage systems) relative to smaller, non-home cage experimental systems and they do not currently offer adaptability.

The first limitation of this study is that it was not possible to directly test the effect of analgesics as described above, as morphine would likely decrease wheel running and weaker analgesics have poor efficacy in models of CIBP [31]. Secondly, there was no difference observed in horizontal home cage activity and use of this system only for wheel running may not be a cost-effective.

This study demonstrates that DVCs® can be an exciting tool to assess pain-like behaviour in different models of pain-like development. It facilitates the simultaneous study of two measures of spontaneous pain-like behaviour over a long period, while reducing the introduction of confounding factors.

To conclude, this is the first study that uses DVC® home cage activity and wheel running to assess pain-like behaviour, here in a model of breast CIBP. This method can be implemented to assess characteristics of other mouse models of painful diseases and perhaps test novel analgesics.

## Acknowledgement

Chelsea Hopkins and Anne-Marie Heegaard received funding for this project from the European Union’s Horizon 2020 research and innovation programme under the Marie Skłodowska-Curie grant agreement No 814244.

